# Centromeres of the yeast *Komagataella phaffii (Pichia pastoris)* have a simple inverted-repeat structure

**DOI:** 10.1101/056382

**Authors:** Aisling Y. Coughlan, Sara J. Hanson, Kevin P. Byrne, Kenneth H. Wolfe

## Abstract

Centromere organization has evolved dramatically in one clade of fungi, the Saccharomycotina. These yeasts have lost the ability to make normal eukaryotic heterochromatin with histone H3K9 methylation, which is a major component of pericentromeric regions in other eukaryotes. Following this loss, several different types of centromere emerged, including two types of sequence-defined ("point") centromeres, and the epigenetically-defined "small regional" centromeres of Candida albicans. Here we report that centromeres of the methylotrophic yeast *Komagataella phaffii* (formerly called *Pichia pastoris*) are structurally-defined. Each of its four centromeres consists of a 2-kb inverted repeat (IR) flanking a 1-kb central core (*mid*) region. The four centromeres are unrelated in sequence. CenH3 (Cse4) binds strongly to the cores, with a decreasing gradient along the IRs. This mode of organization resembles *Schizosaccharomyces pombe* centromeres but is much more compact and lacks the extensive flanking heterochromatic otr repeats. Different isolates of *K. phaffii* show polymorphism for the orientation of the mid regions, due to recombination in the IRs. *CEN4* is located within a 138-kb region that changes orientation during mating-type switching, but switching does not induce recombination of centromeric IRs. The existing genetic toolbox for *K. phaffii* should facilitate analysis of the relationship between the IRs and the establishment and maintenance of centromeres in this species.

## Introduction

Centromeres are the point of connection between the genome and the cytoskeleton during cell division. In most eukaryotes the centromere has a core region that is wrapped around nucleosomes containing the centromeric variant of histone H3 (called CenH3, or Cse4 in yeasts). A large protein complex, the kinetochore, connects these nucleosomes to the spindle microtubules that move the chromosomes during mitosis and meiosis. In most eukaryotes, the core region of the centromere is flanked by extensive arrays of heterochromatin that are repetitive in sequence and largely transcriptionally inert (Pluta et al. 1995; Volpe et al. 2002; Chan and Wong 2012; Hall et al. 2012). The heterochromatin has a characteristic modification, di- or tri-methylation of lysine 9 of histone H3 (H3K9me2/3) that makes it compact. This typical eukaryotic organization is seen in most fungi including basidiomycetes (phylum Basidiomycota), filamentous ascomycetes (subphylum Pezizomycotina) such as *Neurospora crassa*, and ascomycete yeasts related to *Schizosaccharomyces pombe* (subphylum Taphrinomycotina) (Allshire 2004; Smith et al. 2012; Janbon et al. 2014). However, an early ancestor of the subphylum Saccharomycotina lost the enzymatic machinery (Swi6/Clr4/Epe1) for making and maintaining H3K9me2/3 heterochromatin (Malik and Henikoff 2009; Allshire and Ekwall 2015; Riley et al. 2016). Yeasts that are descended from this ancestor have centromeres that are very different from other eukaryotes, being smaller and also very variable among species (summarized in Fig. 1). Their centromeres are usually categorized into two types, ‘point centromeres’ and ‘small regional centromeres’ (Roy and Sanyal 2011), whereas the term ‘large regional centromeres’ is used for those present in H3K9me2/3-containing fungi and other eukaryotes.

**Figure 1.**
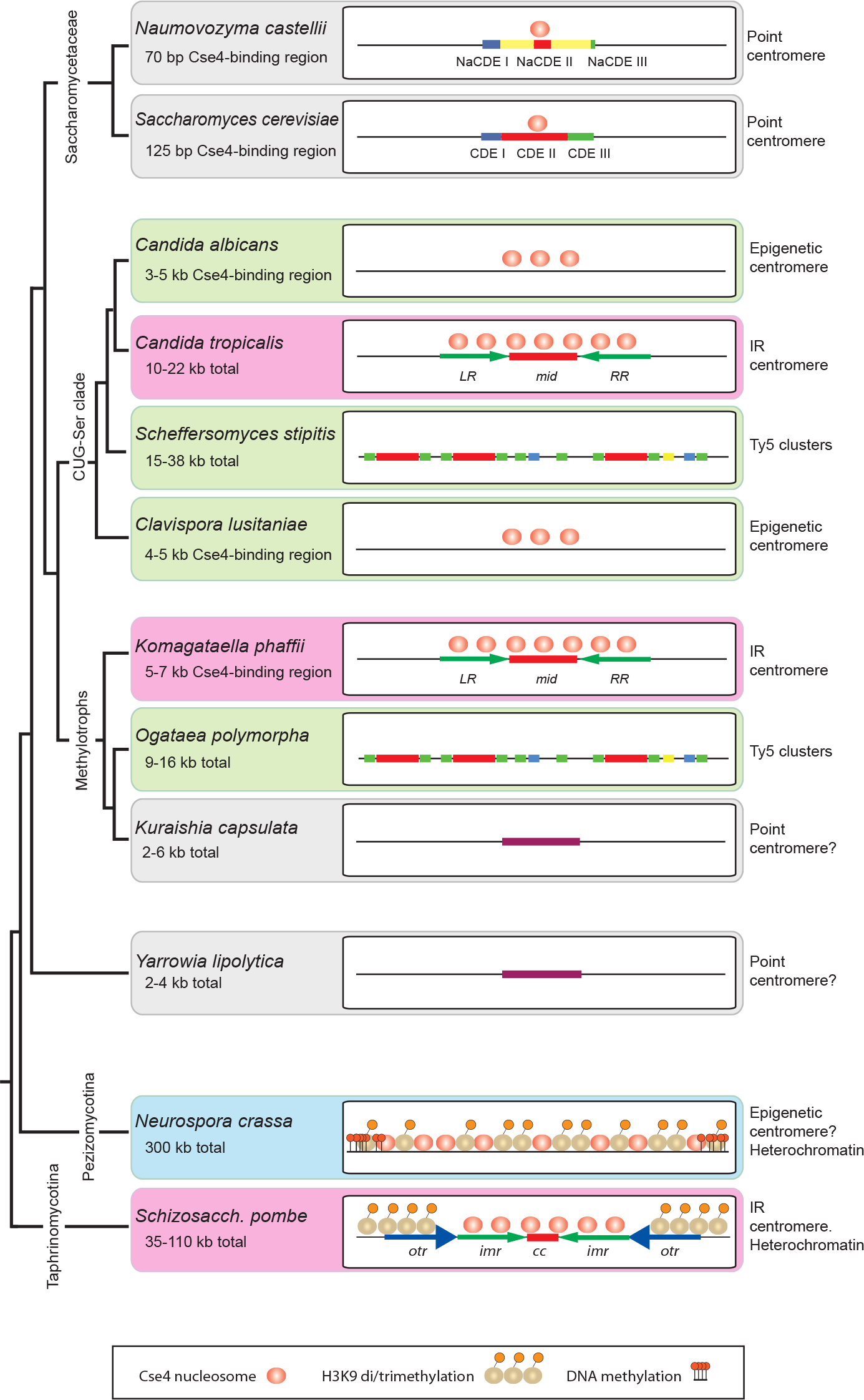
Heterogeneity of ascomycete fungal centromere organization. Schematic (not to scale) showing the structure and epigenetic modification of centromeres across clades of ascomycete fungi. Pink backgrounds indicate IR-containing centromeres; green backgrounds indicate other types of 'small regional’ centromere. Cse4 nucleosome occupancy is unknown for some species.

All studied species of the yeast family Saccharomycetaceae have point centromeres. As first identified in *Saccharomyces cerevisiae* (Fitzgerald-Hayes et al. 1982; Hieter et al. 1985; Hegemann and Fleig 1993), point centromeres are small and have a clear consensus sequence consisting of two short sequence motifs (CDE I and CDE III) to which kinetochore proteins bind, separated by an A+T-rich region (CDE II). Mutations in the conserved (CDE) motifs of point centromeres can result in chromosome instability, and plasmids containing a cloned *CEN* sequence are mitotically stable (Hegemann and Fleig 1993; Kobayashi et al. 2015). In other words, a point centromere’s sequence is necessary and sufficient for centromeric function. Each point centromere of *S. cerevisiae* is only 125 bp long and binds only a single CenH3-containing nucleosome (Henikoff and Henikoff 2012). As well as being well conserved across all *S. cerevisiae* chromosomes (Fleig et al. 1995), the CDE I – CDE III consensus is also well conserved among most species of the family Saccharomycetaceae (Gordon et al. 2011) but a notable exception occurs in the genus *Naumovozyma* (Kobayashi et al. 2015). Although *Naumovozyma* lies within the family Saccharomycetaceae (Fig. 1), and therefore originally had point centromeres similar to those of S. *cerevisiae*, the two examined species in this genus (*N. castellii* and *N. dairenensis*) now have unorthodox point centromeres with a consensus sequence completely unlike that in the rest of the family. Since the *Naumovozyma* centromeres are also not at absolutely conserved chromosomal locations relative to centromeres in other Saccharomycetaceae, it is probable that this new type of point centromere invaded *Naumovozyma* genomes and displaced the old type (Kobayashi et al. 2015).

The CUG-Ser clade shows wide variation in centromere structures (Fig. 1). ‘Small regional’ centromeres were first identified as Cse4-binding regions in *Candida albicans* and its close relative *C. dubliniensis.* These centromeres show no consensus sequence or structural features (Sanyal et al. 2004; Padmanabhan et al. 2008). They range from 4–18 kb of gene-free DNA, and are thus intermediate in size between point centromeres and ‘large regional’ centromeres. About 3–5 kb of each *C. albicans* centromere is occupied by Cse4 (Sanyal et al. 2004; Roy and Sanyal 2011). The sequence of a *C. albicans* centromere appears to be neither necessary nor sufficient for centromeric function. It is not necessary, because if a centromere is deleted a neocentromere can form spontaneously elsewhere on the chromosome (Ketel et al. 2009; Thakur and Sanyal 2013). It is not sufficient, because plasmids or chromosome fragments containing *CEN* regions do not show mitotic stability when transformed into *C. albicans* (Baum et al. 2006). Thus, the mechanism that has kept centromeres at a conserved location, both among strains of *C. albicans* and between *C. albicans* and *C. dubliniensis,* is not well understood (Padmanabhan et al. 2008; Thakur and Sanyal 2013). *C. lusitaniae* has similarly nonrepetitive ‘small regional’ centromeres, located in deep troughs of G+C content, with 4–5 kb occupied by Cse4 (Kapoor et al. 2015). However *C. tropicalis* was recently shown to have centromeres containing a central 2–10 kb region bound by Cse4, flanked by 3–6 kb inverted repeat (IR) structures (Chatterjee et al. 2016). Also in the CUG-Ser clade, the putative centromere regions of *Debaryomyces hansenii* and *Scheffersomyces stipitis* contain large clusters of Ty5-like retrotransposon elements within which the precise location and structure of the centromere remains unknown (Lynch et al. 2010).

The Saccharomycetaceae, CUG-Ser and methylotroph clades are the three major clades of Saccharomycotina (Fig. 1) (Riley et al. 2016). Outside these but still in subphylum Saccharomycotina, *Yarrowia lipolytica* has small centromeres with a weak dyad consensus sequence (Vernis et al. 2001; Yamane et al. 2008), located in G+C troughs (Lynch et al. 2010). These centromeres lie in 2–4 kb intergenic regions and are often described as point centromeres, although the consensus sequence is much more poorly conserved among chromosomes than that in Saccharomycetaceae.

The Pezizomycotina and Taphrinomycotina have ‘large regional’ centromeres with H3K9me2/3 heterochromatin, resembling typical eukaryotic centromeres. The best-characterized species in these subphyla are *Schizosaccharomyces pombe* and *Neurospora crassa.* The centromeres of S. *pombe’s* three chromosomes are 35–110 kb and contain two types of sequence repeat (Takahashi et al. 1992; Baum et al. 1994; Wood et al. 2002; Allshire and Ekwall 2015). Each centromere has a Cse4-binding domain of about 15 kb comprising a non-repetitive central core (cc) flanked by two inverted copies of an innermost repeat *(imr)* sequence which is chromosome-specific (Fig. 1). The pericentromeric regions, marked by H3K9 methylation, consist of 1–9 tandem copies of an outer repeat *(otr)* sequence, arranged in opposite orientations on the two chromosome arms and therefore extending the overall inverted repeat organization of the whole centromere. Whereas the *imr* repeats are different on each chromosome, all the *otr* repeats are similar. Variation in the number of *otr* repeats is the main cause of size difference among the S. *pombe* centromeres. *N. crassa* centromeres consist of 150–300 kb of A+T-rich DNA, rich in heterogeneous repeat sequences and inactivated transposons (Smith et al. 2011). Unlike *S. pombe,* they contain no large IR structures (Roy and Sanyal 2011). H3K9 trimethylation is found throughout the entire *N. crassa* centromeric region, and is required for appropriate CenH3 occupancy (Smith et al. 2011).

In this study, we characterize the centromeres of *Komagataella phaffii,* a haploid species in the methylotrophs clade (Fig. 1). The only previous studies of methylotroph centromeres have been in *Kuraishia capsulata* which has a 200 bp conserved consensus sequence on 5 of its 7 chromosomes (Morales et al. 2013), and *Ogataea (Hansenula) polymorpha* which has Ty5-like retrotransposon clusters without other obvious conservation of centromere sequence or structure (Ravin et al. 2013; Hanson et al. 2014). *K. phaffii* is better known by an obsolete name, *Pichia pastoris*, and is widely used in biotechnology (Kurtzman 2009; Dikicioglu et al. 2014). *K. phaffii* has a compact genome of only 9 Mb and four chromosomes, with a ribosomal DNA array located at one end of each chromosome, and no known transposable elements (De Schutter et al. 2009; Kuberl et al. 2011). Its centromeres were recently mapped with low resolution by a Hi-C method (Varoquaux et al. 2015). Here, we localize the centromeres with high-resolution ChIP-seq and identify an IR structure at each *K. phaffii* centromere, reminiscent of S. *pombe* and *C. tropicalis* but with some significant differences. Thus, following the loss of H3K9me2/3, centromere structure has continued to evolve in diverse directions within each clade of Saccharomycotina, as well as between the clades.

## Results

### One transcription coldspot per chromosome in K. phaffii

An extensive RNAseq dataset for transcriptomes of glycerol-grown *K. phaffii* cells was published by Liang et al. (2012). We mapped these reads to the *K. phaffii* genome, which in general is very compact with little noncoding DNA (Kuberl et al. 2011), and noticed that there is one large non-transcribed region on each chromosome (Figs. 2,3). These non-transcribed regions are 6–9 kb long.

Each non-transcribed region contains a simple IR structure, with two almost identical copies of a 2 kb sequence separated by about 1 kb (Fig. 3; Table 1). We hypothesized that these IR-containing regions could be centromeres. They are reasonably close to the previous estimates of centromere locations by Hi-C (Varoquaux et al. 2015), ranging from 0–19 kb away (Table 1). Independently, Sturmburger *et al.* (2016) also proposed these IR regions as candidate centromeres.

**Table I.** Chromosomal coordinates and sizes of the components of the centromeres of K. phaffii strain CBS7435.

| Chr. | IR<br>length<br>(bp) | Diffs.<sup>a</sup> | *mid*<br>length<br>(bp) | Total<br>*CEN*<br>length<br>(bp) | Left Repeat (*LR*)<br>coordinates<sup>b</sup> | Right Repeat (*RR*)<br>coordinates<sup>b</sup> | Varoquaux *et al.*<br>20 kb call<sup>c</sup> 40 kb call<sup>c</sup> |  |
| --- | --- | --- | --- | --- | --- | --- | --- | --- |
| 1 | 2012 | 1 | 1330 | 5354 | 1401537 - 1403548 | 1404879 - 1406890 | 1408922 | 1404619 |
| 2 | 2699 | 1 | 1257 | 6655 | 843845 - 846543 | 847801 - 850499 | 838077 | 837858 |
| 3 | 2650 | 1 | 883 | 6182 | 34396 - 37045 | 37929 - 40578 | 18617 | 35592 |
| 4 | 2559 | 0 | 1111 | 6229 | 78869 - 81427 | 82539 - 85097 | 76386 | 69315 |

<sup>a</sup> Number of nucleotide differences between the *LR* and *RR*.
<sup>b</sup> CBS7435 genome coordinates of Kuberl *et al.* (2011). Accession numbers FR839628.1 to FR839631.1.
<sup>c</sup> Hi-C estimates of centromere position by Varoquaux *et al.* (2015) using two methods. GS115 coordinates were converted to CBS7435 coordinates.

### Cse4 ChIP-seq confirms IR regions as centromeres

Chromatin immunoprecipitation is a technique that has been used widely to locate centromeres in other yeasts (Sanyal et al. 2004; Lefrancois et al. 2013; Hanson et al. 2014; Kobayashi et al. 2015; Chatterjee et al. 2016). To locate the *K. phaffii* centromeres, we used ChIP-seq to identify sequences associated with centromeric nucleosomes containing CenH3 (Cse4). We constructed a strain of *K. phaffii* with a haemagglutinin (3xHA) tag at an internal site (Stoler et al. 1995; Wisniewski et al. 2014) of the endogenous *CSE4* gene (Fig. S1).

We obtained a single robust ChIP-seq signal for a centromere on each of the four chromosomes (Fig. 2). The signal is highest in the non-repetitive central *(mid)* region between the two IRs of each chromosome, and declines along the left and right arms of the IR *(LR* and *RR* in Fig. 3). The region of Cse4 binding coincides almost exactly with the region of no transcription.

**Figure 2.**
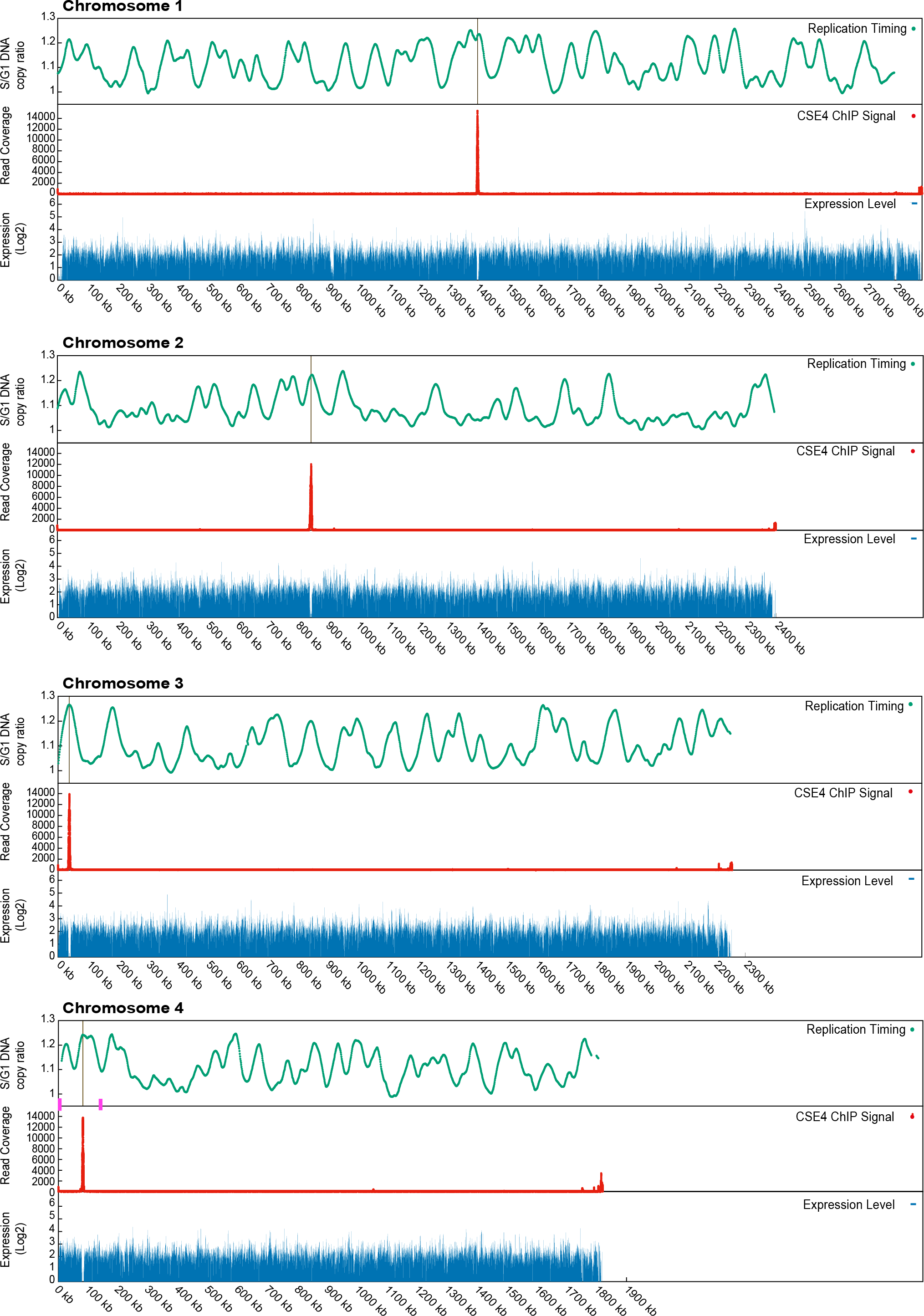
Cse4 occupancy, replication timing and transcription density along the 4 *K. phaffii* chromosomes. Red, ChIP-seq signal from HA-tagged Cse4. The single large peak on each chromosome identifies the centromere. Weaker signals at the right end of each chromosome coincide with ribosomal DNA repeats. Green, replication data from Liachko et al. (2014) re-mapped to the genome of strain CBS7435. Peaks indicate early replicating regions. Blue, transcription data from RNA-seq experiment of Liang et al. (2012) for cultures grown in glycerol. The gaps in transcription at 900 kb and 2800 kb on chromosome 1 are due to the very large gene *MDN1* and a gap (poly-N region) in the genome sequence, respectively. Magenta boxes indicate the positions of the two *MAT* loci on chromosome 4 (Hanson et al. 2014).

**Figure 3.**
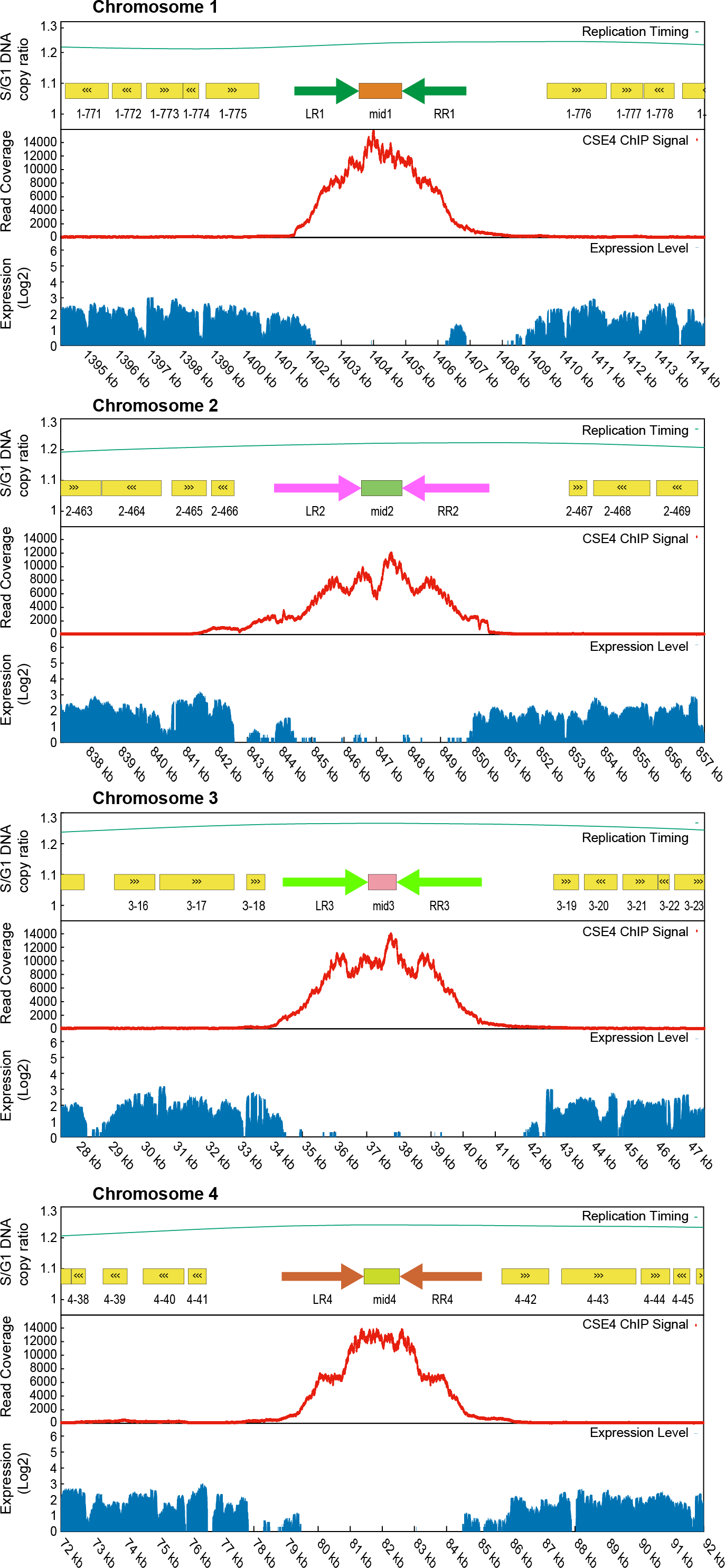
Inverted repeat structure and Cse4 occupancy in centromeric regions. A 20kb region centered on the highest Cse4 peak is shown for each chromosome. Cse4 occupancy, transcription and replication data are as in Fig. 2. Paired arrows show the locations of the IRs flanking each *mid* region. The sequences of *LR* and *RR* are virtually identical on each chromosome, but different between chromosomes; *mid* sequences are different between chromosomes. Yellow boxes indicate annotated protein-coding genes.

The *LR* and *RR* of each centromere are virtually identical to each other (Table 1), but completely unrelated on each chromosome despite being similar in size. The *mid* regions of each centromere are also unrelated. There is, however, a small region of similarity between the *mid* region of *CEN1* and the IRs of *CEN4* which is visible in a dot matrix plot (Fig. S2). The aligned region has 65% sequence identity over 518 bp. Other than this region, we could not find any conserved sequence motifs in the *K. phaffii* centromeres using MEME (Tanaka et al. 2014). The base composition of the centromeres is not exceptional: the *mid* regions range from 37.4–40.7% G+C, and the IRs from 37.3–38.2% G+C; for comparison the whole genome averages 41.1% G+C.

Interestingly, *CEN4* is located within a 138 kb section of chromosome 4 whose orientation becomes inverted during mating-type switching (Hanson et al. 2014). In this process, recombination occurs between two identical 2.6 kb sequences (the ‘outer IR’) that lie in inverted orientations, one near the expressed copy of the *MAT* genes and the other near the silenced copy. The centromere is in the approximate center of the invertible region, 58 kb and 79 kb from the two *MAT* locus outer IRs (Fig. 2).

It is also notable that *CEN3* and *CEN4* are both located close to one end of their chromosomes (37 kb and 82 kb from the telomere, respectively), making the right arm 60 times longer than the left on chromosome 3, and 22 times longer on chromosome 4. In both cases the centromere is at the opposite end of the chromosome from the rDNA array.

### Centromeres are early-replicating

Because cell division can only proceed after the centromeres have replicated and kinetochores have been assembled on them, centromeres tend to be early-replicating in all eukaryotes (McCarroll and Fangman 1988; Kim et al. 2003; Koren et al. 2010). Liachko *et al.* (2014) previously mapped replication origins throughout the *K. phaffii* genome, using a high-throughput sequencing method to compare the copy number of 1-kb windows of the genome in S versus G1 phase of the cell cycle in *K. phaffii* strain GS115. We re-mapped their data to coordinates for strain CBS7435, whose genome is almost identical to GS115 (Kuberl et al. 2011). The four centromeres are all seen to be located at peaks of early replication (Figs. 2,3, green lines), though each chromosome contains multiple similarly early peaks, confirming the result from the lower resolution Hi-C study (Varoquaux et al. 2015). The two *MAT* locus regions on chromosome 4 are both late-replicating (Fig. 2).

### Polymorphic inversions at centromeres

We searched for identical DNA sequence repeats in the *K. phaffii* genome, and found that there are only 7 such repeats larger than 1 kb: the four centromeric IRs, the inner and outer IRs flanking the *MAT* locus (Hanson et al. 2014), and the ribosomal DNA arrays at the right telomere of each chromosome. We previously found that natural isolates of *K. phaffii* are polymorphic for the orientation of the 138-kb region of chromosome 4, due to recombination between the *MAT* locus IRs. Recombination in the 2.6 kb outer IR of the *MAT* locus is induced during mating-type switching, whereas recombination in the 5.7 kb inner IR is not inducible but appears to occur in nature (Hanson et al. 2014).

To test whether the orientation of centromeres is polymorphic due to recombination in their IRs, we used orientation-specific PCR assays to determine the arrangement of each *mid* region relative to the flanking chromosome arms. Among five natural isolates of *K. phaffii* that we tested, the orientations of three centromeres *(CEN1, CEN2* and *CEN4)* are variable (Fig. 4), indicating that ectopic recombination between the centromeric IRs also occurs in nature.

**Figure 4.**
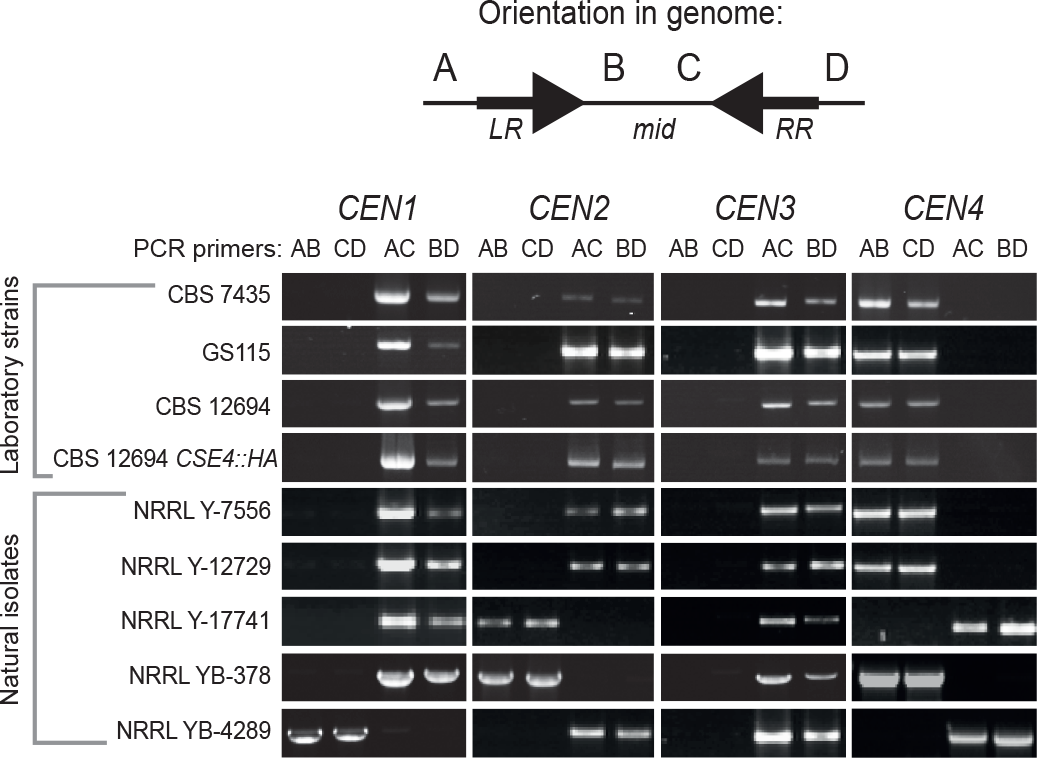
Polymorphism of centromere orientation among *K. phaffii* strains. Genomic DNAs from 4 haploid laboratory strains and 5 natural isolates of *K. phaffii* from the USDA Agricultural Research Service NRRL collection (Kurtzman 2009) were amplified using orientation-specific PCR assays. For each centromere, two primers (B and C) bind to the *mid* region and two (A and D) bind to the chromosome arms outside the IR regions. The 16 primer sequences are listed in Table S1. Amplification with the AB and CD primer pairs for a centromere indicates that its *mid* region is in the same orientation as the reference genome sequence (Kuberl et al. 2011) whereas amplification with the AC and BD pairs indicates inversion of the *mid* region relative to the chromosome arms. PCR products are the size of the IRs (2.0–2.7 kb; Table 1). PCR amplifications used GoTaq (Promega) polymerase for 30 cycles with an annealing temperature of 55°C.

We found that centromere orientation is conserved between the laboratory strain CBS7435 and three derivatives of this strain (Fig. 4). These derivatives are the biotech strain GS115, the *ku70* knockout strain CBS12694 (Naatsaari et al. 2012), and the *ku70 CSE4∷HA* strain we used for the ChIP-seq experiment. However, the PCR-determined orientation of *CEN1, CEN2*, and *CEN3* in all the laboratory strains is opposite to that reported in the genome sequences of CBS7435 (Kuberl et al. 2011) and GS115 (De Schutter et al. 2009), probably due to propagated errors in genome assembly.

Because the centromeric IRs are similar in size to the *MAT* locus IRs, and because *CEN4* is located within the 138–kb region that inverts during mating-type switching, we were curious whether induction of mating-type switching might also induce inversion of some centromeres. We induced mating-type switching by growing strain GS115 (MATalpha) in liquid NaKG media (Hanson et al. 2014), then plating for single colonies on YPD, and isolating genomic DNA from multiple cultures each grown from a single colony. Among these cultures, we identified four switched *MATa* clones by *MAT* locus orientation-specific PCR (Hanson et al. 2014). We then assayed their centromere orientations. However, the orientations of all four centromeres remained unchanged in the four *MATa* clones (Fig. S3).

## Discussion

*K. phaffii, C. tropicalis* and *S. pombe* are the only three fungi known to have centromeres with an IR organization, but there are significant differences among these species. They are compared in dotplots in Figures S2, S4 and S5. All seven centromeres of *C. tropicalis* are similar to one another (>60% sequence identity in both their *mid* and IR regions; Fig. S4) suggesting that they have been homogenized by gene conversion (Chatterjee et al. 2016). In contrast, each centromere of *K. phaffii* has a unique sequence, except for a small patch of gene conversion between *CEN1* and *CEN4* (Fig. S2). Similarly, each *imr-cc* region of *S. pombe* is different, except for a patch of gene conversion (*tm* region) between *cc1* and *cc3* (Fig. S5; Takahashi et al. 1992). The centromeres of *K. phaffii* therefore strongly resemble the structure of the *imr-cc* regions at the heart of *S. pombe* centromeres (and other Schizosaccharomyces species; Rhind et al. 2012) but they lack the additional nested arrays of *otr* repeats that extend the IRs and are similar across all S. *pombe* chromosomes (Fig. S5). In both *K. phaffii* (this study) and *S. pombe* (Steiner et al. 1993), natural isolates are polymorphic for the orientation of the central region relative to the chromosome arms, for at least some centromeres.

In both *K. phaffii* and *S. pombe*, the CenH3 ChIP-seq signal is maximal in the unique region (*mid* or cc) and decays along the flanking IRs *(imr)* (Fig. 3; Thakur et al. 2015). In *C. tropicalis*, Chatterjee *et al.* (2016) reported that Cse4 is located only in the *mid* regions and not on the IRs. However, this conclusion is an artifact of the parameters they used to map the ChIP-seq data with the program Bowtie (Langmead and Salzberg 2012), which caused any reads with multiple matches to the genome to excluded. When we re-mapped their data to the *C. tropicalis* genome allowing up to two matches per read to be reported, we find a decreasing signal along each *C. tropicalis* IR similar to Fig. 3. Therefore, all three species have a similar pattern of CenH3 distribution, which suggests that the IRs may have a function in specifying where CenH3 nucleosomes are placed. A possible evolutionary role of the IRs is to prevent drift of the centromere position onto neighboring genes, affecting their transcription. This is supported by studies in *C. albicans*, which showed that neocentromeres are capable of moving locally along DNA under certain conditions (Ketel et al. 2009).

The major difference between the *S. pombe* centromeres and those of *K. phaffii* and *C. tropicalis* is the presence of the outer repeats *(otr)* and their associated heterochromatin. H3K9 methylation at *otr* is established by bidirectional transcription. The RNAi machinery processes these transcripts to direct the H3K9 methyltransferase (Clr4) to this region of the centromere (Volpe et al. 2002). In contrast, the Saccharomycotina yeasts lack most of the components of this system: there are no outer repeats and no H3K9 methylation proteins (Clr4/Swi6/Epe1) in either *K. phaffii* or *C. tropicalis*, and no RNAi apparatus (Ago1/Dcr1) in *K. phaffii.* Since heterochromatic centromeres are the norm in most eukaryotes, we must presume that the Saccharomycotina IR-containing centromeres are derived from heterochromatic ancestors and have become simplified. What is the function of *otr?* Assays in *S. pombe* found that a partial *otr* repeat and most of the central domain is sufficient for establishing a functional centromere on naive DNA (Baum et al. 1994; Steiner and Clarke 1994; Ngan and Clarke 1997), but *otr* and H3K9me2/3 heterochromatin are not required for maintenance of an already active centromere (Baum et al. 1994; Folco et al. 2008). CenH3 when overexpressed can bind specifically to a central core (cc2) region entirely lacking both *otr* and *imr* (Catania et al. 2015). These results indicate the central cores of *S. pombe* centromeres have some feature that gives them an innate ability to attract CenH3. Once CenH3 chromatin is established, it is capable of self-propagation and thus is the primary epigenetic mark for kinetochore formation.

The diversity of eukaryotic centromere structures is a consequence of rapid coevolution between centromere DNA and the kinetochore proteins (Padmanabhan et al. 2008; Malik and Henikoff 2009; Bensasson 2011; Kobayashi et al. 2015). Ascomycete centromeres have more structural diversity than can be conveyed by a simple binary distinction between point and regional centromeres. We suggest that it might instead be useful to consider yeast centromeres in terms of whether their CenH3-binding region is sequence-defined, structure-defined or epigenetically-defined. Point centromeres are sequence-defined, meaning that a specific DNA sequence is required for them to function. Structure-defined centromeres require a particular arrangement of repeat sequences in the CenH3-bound region, such as the IRs in *K. phaffii, C. tropicalis* and *S. pombe*, but the actual sequence of the repeating units may not be important. Epigenetically-defined centromeres are those such as in *C. albicans* and *C. lusitaniae*, where the only factor determining where new CenH3 nucleosomes are deposited appears to be the presence of CenH3 in the previous generation. It is unclear whether species with Ty5-like retrotransposon clusters at their centromeres constitute a different (fourth) group; detailed CenH3 mapping has not been carried out in any yeasts with this structure, and it is possible that the kinetochore is localized to specific discrete sites and not across the whole retrotransposon cluster. The IR-containing centromere structure may be ancestral in ascomycetes, because it is present in two subphyla of Ascomycota and two families of Saccharomycotina, in which case the epigenetic and Ty5- cluster categories would represent degenerated forms of IR-containing centromeres. In this regard, it is interesting that one centromere *(CEN5)* of *C. albicans* has a large IR (Sanyal et al. 2004; Chatterjee et al. 2016) and that neocentromeres in this species frequently contain repeats (Ketel et al. 2009).

Further experiments will be needed to identify the components required for IR centromere establishment and maintenance in *K. phaffii*, and to determine the role of the IRs. It will be of interest to know what happens if the IRs are deleted, and whether the current IRs can be replaced by other pairs of identical sequences. In *C. tropicalis*, the IRs were found to enhance the activity of *CEN8* in a plasmid stability assay compared to a *mid8* sequence alone, and the IRs of *CEN8* could not be replaced by several alternative sequences that were tested (Chatterjee et al. 2016). *K. phaffii* has several features that could make it an attractive system for further dissection of IR-containing centromeres: it has only four chromosomes; their centromere sequences are all different; and powerful tools for genetic analysis and genome manipulation have already been developed for this species (Naatsaari et al. 2012; Sturmberger et al. 2016; Weninger et al. 2016).

## Acknowledgments

This study was supported by Science Foundation Ireland (13/IA/1910) and the European Research Council (268893). High-throughput sequencing services were performed at the University of Missouri DNA Core Facility.

## Methods

<u>Tagging the endogenous *K. phaffii CSE4* gene.</u> We synthesized a DNA fragment containing the *K. phaffii* CBS7435 *CSE4* gene with a 3xHA (haemagglutinin) tag inserted between amino acids 41 and 42 (Fig. S1). We chose this tagging site based on a multiple alignment of Cse4 proteins that had previously been tagged successfully, after our initial attempt to tag *CSE4* at the C-terminus yielded no viable clones. Fusion PCR was used to fuse the synthetic fragment to a kanamycin resistance *(kanMX)* marker and a short region downstream of the endogenous *K. phaffii CSE4* gene (Fig. S1). The cassette was transformed into a *ku70* mutant strain (strain CBS12694, a derivative of CBS7435; Naatsaari et al. 2012) by electroporation (Lin-Cereghino et al. 2005). Integrants were selected on YPD containing 200 μg/ml G418. Colonies were screened for correct integration by PCR and verified by sequencing.

<u>ChIP-seq.</u> Chromatin immunoprecipitation followed Hanson *et al.* (2014). Yeast cells were cultured in 200 ml YPD to log phase and then crosslinked with 1% formaldehyde at room temperature. Crosslinking was stopped by addition of 2.5 M glycine. Crosslinked cells were washed with TBS (100 mM Tris-HCl, pH 7.5, 150 mM NaCl) and resuspended in FA lysis buffer (50 mM HEPES, pH 7.5, 150 mM NaCl, 1 mM EDTA, 1% v/v Triton X-100, 0.1% w/v sodium deoxycholate, 0.1% w/v SDS) containing 1 mM PMSF. Cells were lysed with glass beads and chromatin was fragmented by sonication with a Bioruptor Standard (Diagenode). Immunoprecipitation of chromatin fragments was performed with EZview Red Anti-HA Affinity Gel (Sigma-Aldrich) or mouse IgG1 (Cell Signaling Technology) as isotype control. After washes, bound DNA was eluted with HA peptide (Sigma-Aldrich), crosslinks were reversed, and the samples were purified by phenol-chloroform extraction before sequencing. Unpaired Illumina ChIP-seq reads (51 bp) were mapped to the 2011 version of the *K. phaffii* CBS7435 genome (Kuberl et al. 2011) including the mitochondrial genome, using BWA v0.7.9a–r786 (aln/samse algorithm) with default parameters (Li and Durbin 2009). Samtools v0.1.12a (r862) was used to create sorted and indexed BAM files of the results. Bedtools v2.19.0 was used to create genome coverage Bedgraph files to produce Figures 2 and 3. We also used Bowtie (Langmead and Salzberg 2012) to re-map the *C. tropicalis* ChIP-seq data with parameters –`m`1 (to replicate Chatterjee et al. 2016) and –`m`2 (to permit reads with up to two matches to be reported).

<u>Replication data:</u> We converted the normalized replication ratio data from Table S5 of Liachko et al. (2014), which were reported for 1 -kb bins in the genome sequence of *K. phaffii* strain GS115, into equivalent coordinates for strain CBS7435. GS115 is a laboratory derivative of CBS7435 and therefore has an almost identical genome, but the published genome sequences of GS115 (De Schutter et al. 2009) and CBS7435 (Kuberl et al. 2011) differ in some details including the orientations of chromosomes 2, 3 and 4, as well as the locations of gaps and the completeness of telomeric regions. To map the replication data, we aligned the sequences of each chromosome between the two strains using MUMMER (Delcher et al. 1999) and calculated a nucleotide-to-nucleotide coordinate conversion table. In Figures 2 and 3, the normalized replication ratio for each 1 -kb bin in GS115 is plotted at the CBS7435 coordinate corresponding to the center of that bin.

---

**Figure S1.**
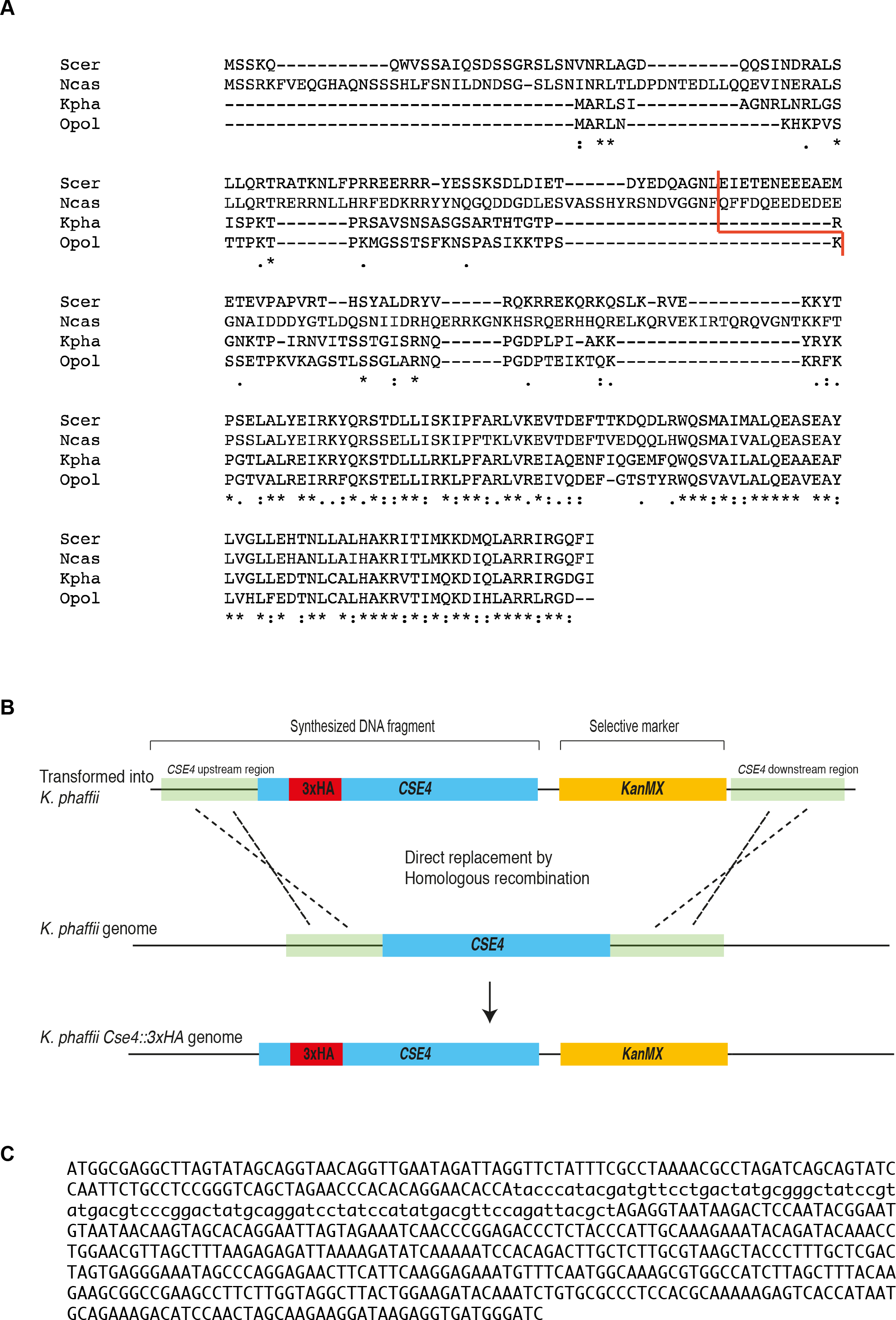
Construction of 3xHA-tagged *CSE4*. (A) Multiple sequence alignment of Cse4 proteins. The red line shows the site of HA tag insertion previously used for ChIP in *S. cerevisiae* (Scer; Stoler et al. 1995), *Naumovozyma castellii* (Ncas; Kobayashi et al. 2014), *Ogataea polymorpha* (Opol; Hanson et al. 2014) and *K. phaffii* (Kpha; this study). (B) Strategy for tagging *K. phaffii CSE4*. The synthetic DNA fragment, *KanMX* marker and *CSE4* downstream DNA were joined by fusion PCR. The entire cassette was then transformed into *K. phaffii* CBS12964 to replace the endogenous *CSE4* gene by homologous recombination. (C) Sequence of the synthetic DNA. The 3xHA tag is in lowercase.

**Figure S2.**
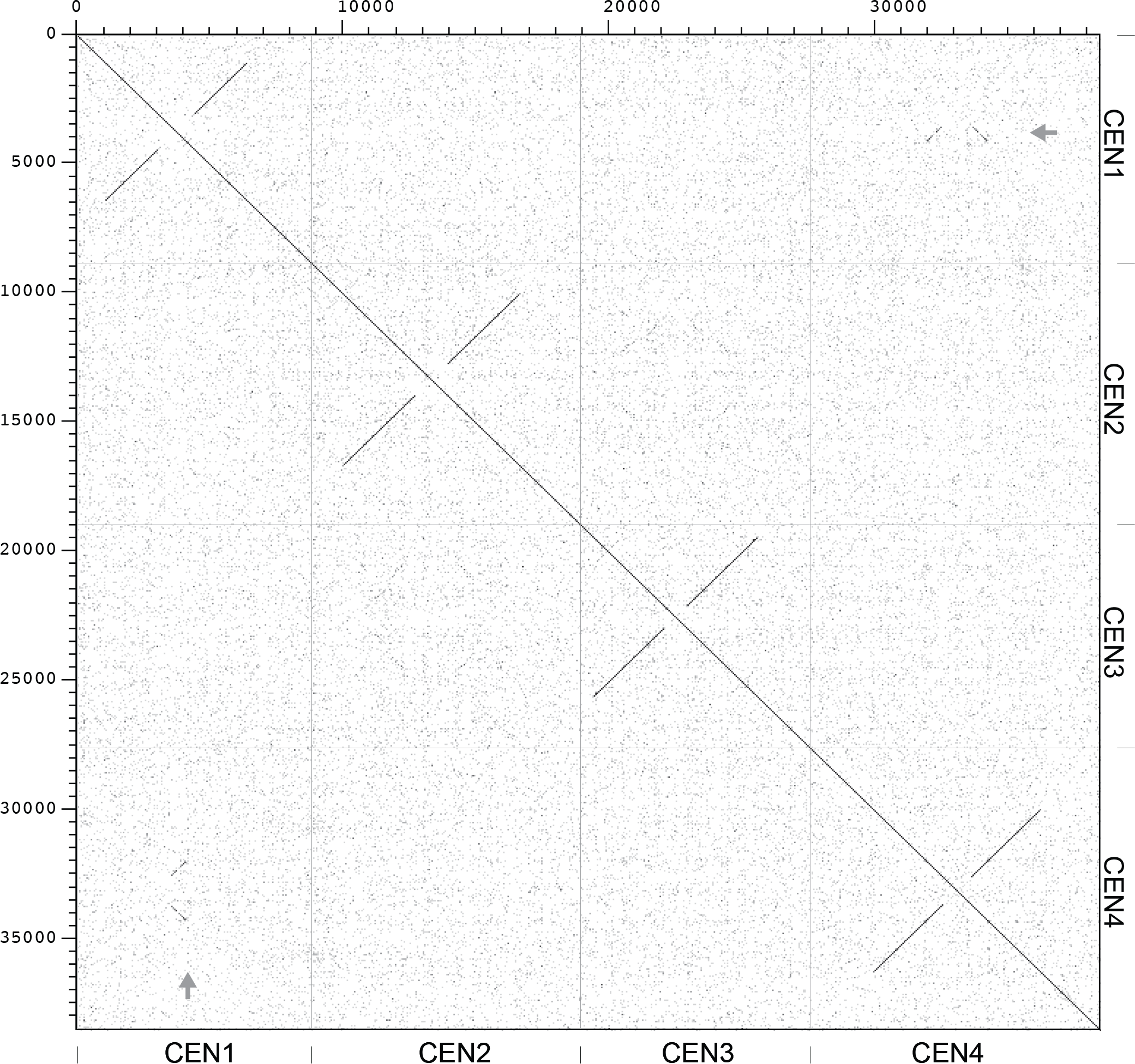
Dot matrix plot of the four *K. phaffii* centromeres compared to each other. Regions of approximately 10 kb around each centromere were concatenated and compared. The arrows indicate a small similarity between *CEN1* and *CEN4*. The plot was constructed with Dotter (Sonnhammer and Durbin 1995), with the Greyramp parameters set to 40 (min.) / 100 (max.).

**Figure S3.**
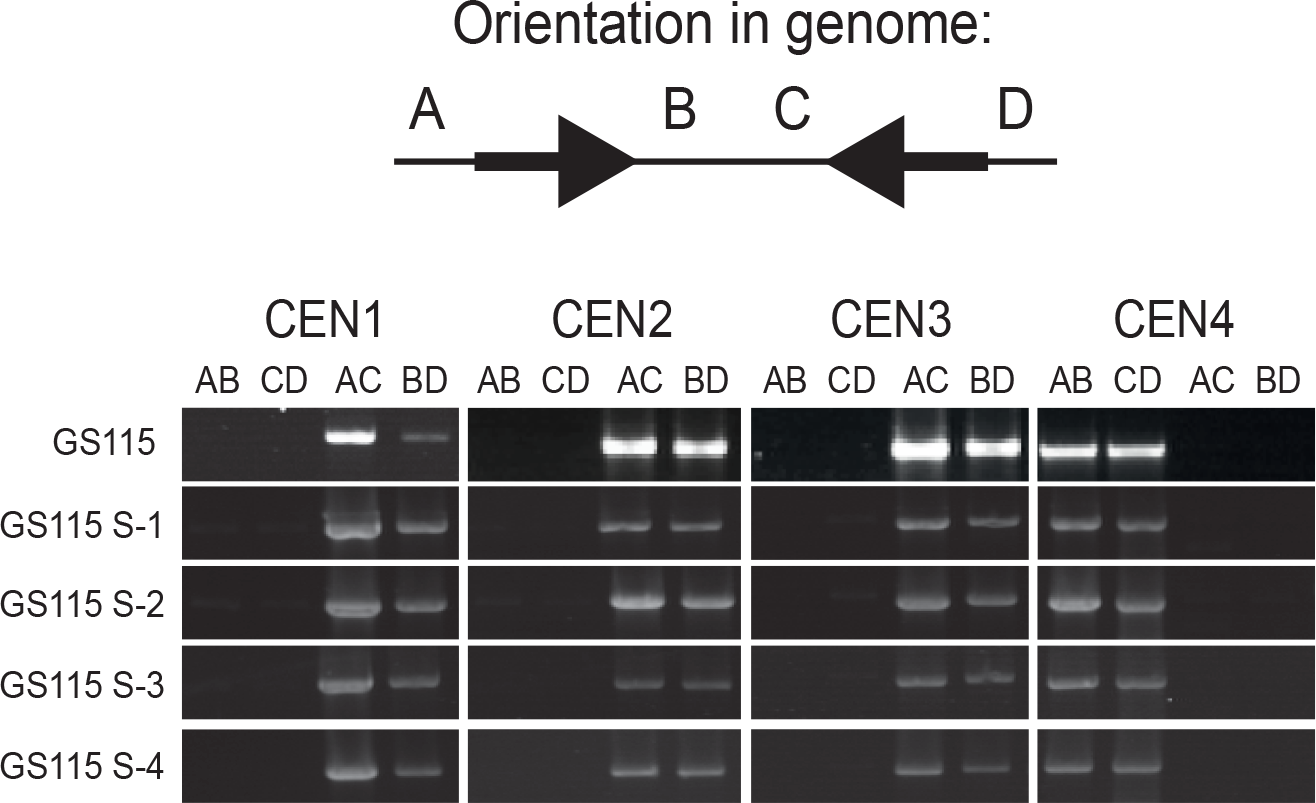
Mating-type switching does not induce recombination at centromeric IRs. GS115 S-1 to S-4 are four *MATa* clones induced by mating-type switching of a GS115 *MAT*alpha strain. Centromere orientation-specific PCR was carried out with primers A-D for each centromere as in Figure 4.

**Figure S4.**
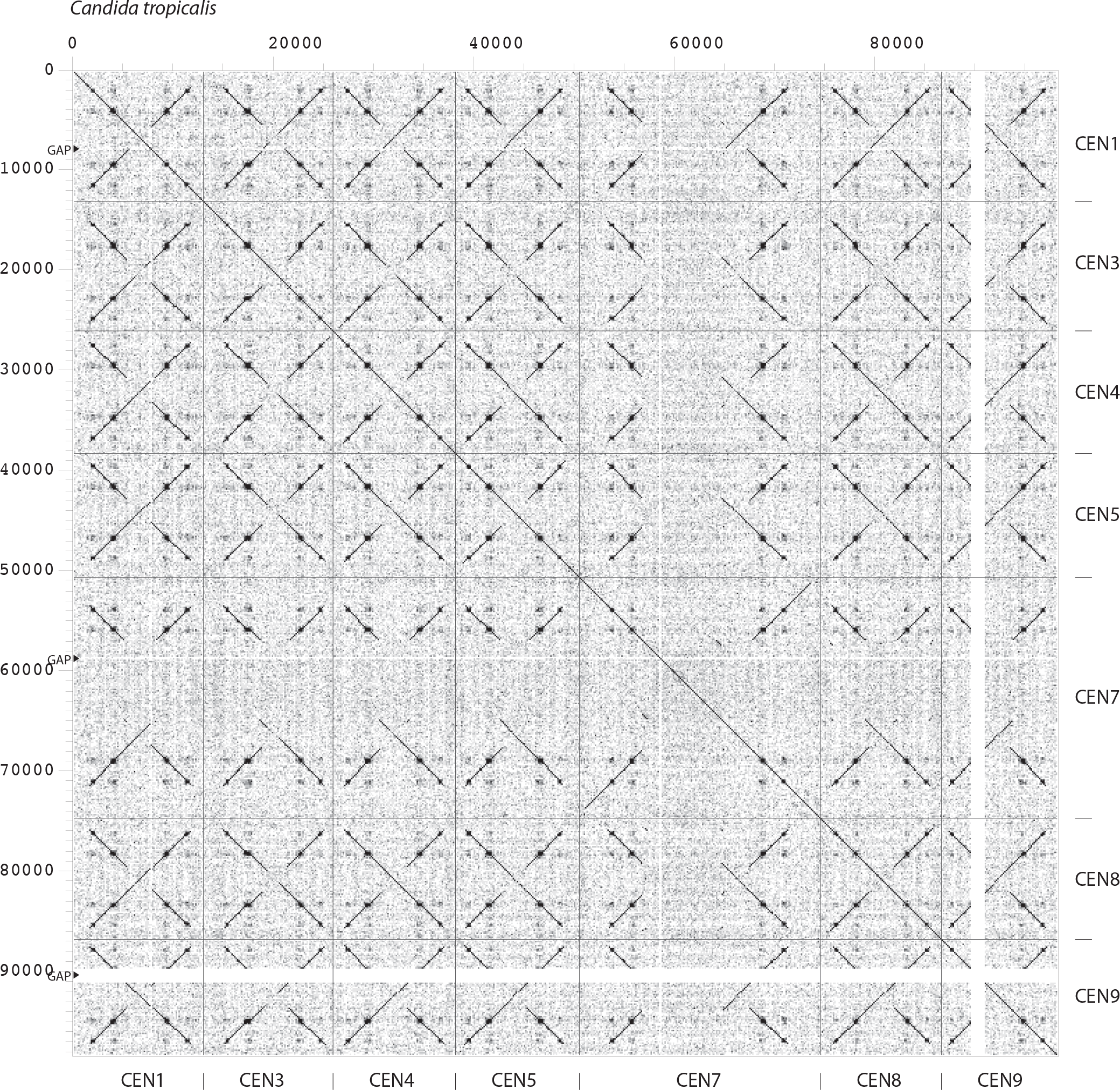
Dot matrix plot of the seven *C. tropicalis* centromeres compared to each other. Parameters are identical to Figure S2. *C. tropicalis* sequence data is from Butler et al. (2009) with centromeres identified by Chatterjee et al. (2016).

**Figure S5.**
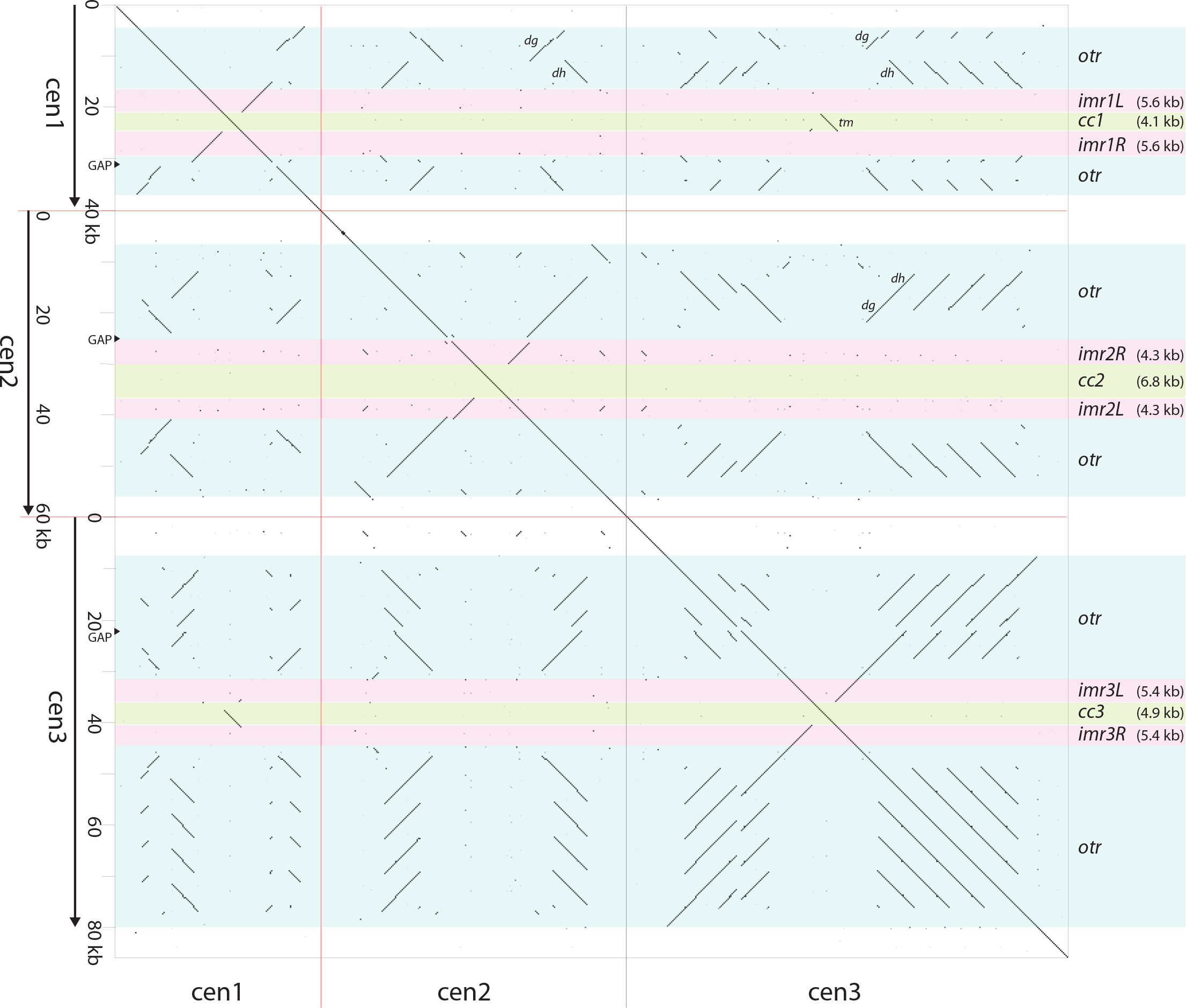
Dot matrix plot of the three *S. pombe* centromeres compared to each other. Red lines indicate points of concatenation between chromosomes. The plot was constructed with Dotter (Sonnhammer and Durbin 1995), with the Greyramp parameters set to 100 (min.) / 150 (max.) to accommodate the larger scale of this figure compared to Figures S2 and S4. The diagram was constructed using the reference genome sequence of *S. pombe* which lacks some copies of *otr* units as indicated by the word GAP on the Y-axis (see Fig. 1 of Wood et al. 2002). The *dg* and *dh* components of the *otr* are marked; *dh* has a different orientation on *cen1* compared to *cen2* and *cen3.* The *tm* region of similarity between *cc1* and *cc3* is also marked.

**Table S1.**
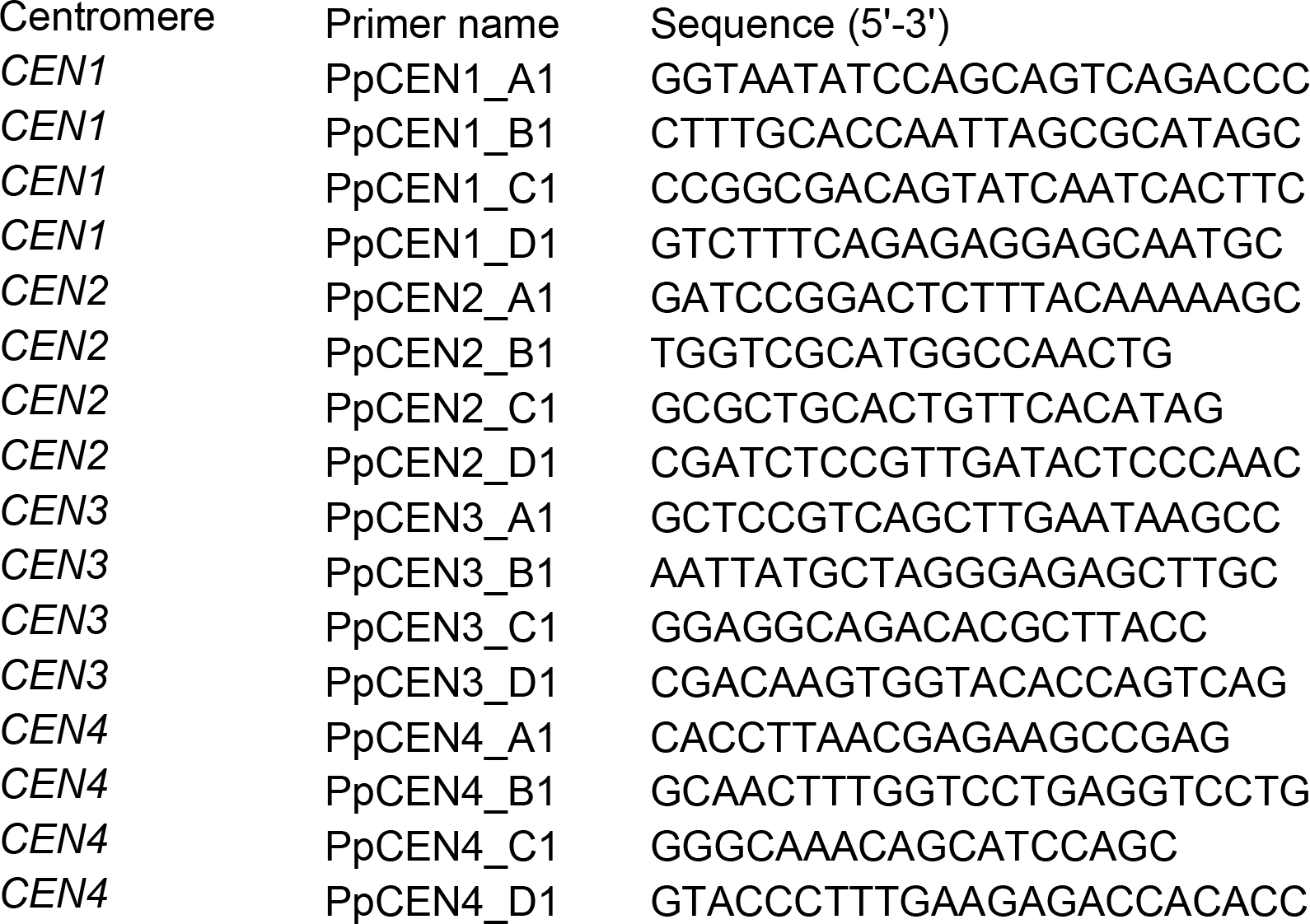
Sequences of primers used for centromere orientation-specific PCR.

